# Mechanism through which retrocyclin targets flavivirus multiplication

**DOI:** 10.1101/2020.12.15.422996

**Authors:** Xiaoying Jia, Jiao Guo, Weirong Yuan, Lingling Sun, Yang Liu, Minmin Zhou, Gengfu Xiao, Wuyuan Lu, Alfredo Garzino-Demo, Wei Wang

## Abstract

Currently, there are no approved drugs for the treatment of flavivirus infection. Accordingly, we tested the inhibitory effects of the novel θ-defensin retrocyclin-101 (RC-101) against flavivirus infection, and investigated the mechanism underlying the potential inhibitory effects. First, RC-101 robustly inhibited both Japanese encephalitis virus (JEV) and Zika virus (ZIKV) infections. RC-101 exerted inhibitory effects on the entry and replication stages. Results also indicated that the non-structural protein NS2B-NS3 serine protease might serve as a potential viral target. Further, RC-101 inhibited protease activity at the micromolar level. We also demonstrated that with respect to the glycoprotein E protein of flavivirus, the DE loop of domain III, which is the receptor-binding domain of the E protein, might serve as another viral target of RC-101. Moreover, a JEV DE mutant exhibited resistance to RC-101, which was associated with deceased binding affinity of RC-101 to DIII. These findings provide a basis for the development of RC-101 as a potential candidate for the treatment of flavivirus infection.

**Importance:** RC has been reported to have a broad-spectrum antimicrobial activity. In this study, we firstly report that RC-101 could inhibit ZIKV and JEV infections. Moreover, both the NS2B-NS3 serine protease and the DE loop in the E glycoprotein might serve as the viral targets of RC-101.

## Introduction

Flaviviruses are taxonomically classified in the genus *Flavivirus* and family Flaviviridae. These viruses include more than 70 different pathogens and are transmitted mostly by arthropods. Emerging and re-emerging flaviviruses, such as Zika virus (ZIKV), Japanese encephalitis virus (JEV), dengue virus (DENV), West Nile virus (WNV), and yellow fever virus, cause public health problems worldwide (1). Flaviviruses contain an approximately 11-kb positive-stranded RNA genome that encodes three structural proteins, including the capsid (C), membrane (premembrane [prM] and membrane [M]), and envelope (E), as well as seven nonstructural proteins (NS1, NS2A, NS2B, NS3, NS4A, NS4B, and NS5) (2). The envelope glycoprotein (E) is responsible for receptor binding and membrane fusion and thus plays essential roles in virus entry. E proteins exist as homodimers on the surface of the virus. Among the three domains of the E protein, domain I (DI) connects the DII and DIII domains, and DII contains fusion polypeptides that facilitate membrane fusion, whereas DIII has been proposed to act as the receptor binding region (3-5). It has been reported that several key residues, such as the glycosylation site N154 and the DE loop (T_363_SSAN_367_) are responsible for receptor binding (6, 7), whereas H144 and H319 are thought to play critical roles in DI and DIII interactions (8). Moreover, Q258 located in DII and T410 located in the stem are indispensable for low pH-triggered conformational changes, in which the stem region undergoes zippering along with DII, thus leading to the post-fusion conformation and membrane fusion (9-11). As it envelops the surface of the virion, the E protein is the natural target for antibodies and the design of entry inhibitors to prevent receptor-binding and membrane fusion (4, 9, 12, 13). Likewise, viral proteases such as NS2B-NS3 protease-helicase and the NS5 RNA-dependent RNA polymerase represent attractive drug targets in an attempt to identify replication inhibitors (14, 15).

Retrocyclin (RC) is an artificially humanized θ-defensin that has been reported to possess broad antimicrobial activity (16-21). RC-101 contains 18 residues including three disulfide bonds and four positively charged residues (Fig. 1A and B), which confers high binding affinity to glycosylated proteins, such as HIV gp120 (22), influenza hemagglutinin (23), and HSV1/2 glycoprotein (24), thus preventing virus entry. Additionally, some viral proteases with negatively charged surfaces might serve as targets for RC (20). In this study, we tested the inhibitory effect of RC-101 against flavivirus infection. As flaviviruses possess only one conserved N-linked glycan on the E protein (25), whether RC exerted the inhibitory effect against flavivirus entry by targeting the glycan chain was tested in this study. Meanwhile, we determined that RC-101 could also inhibit flavivirus replication by blocking the NS2B-NS3 serine protease.

**Fig. 1.**
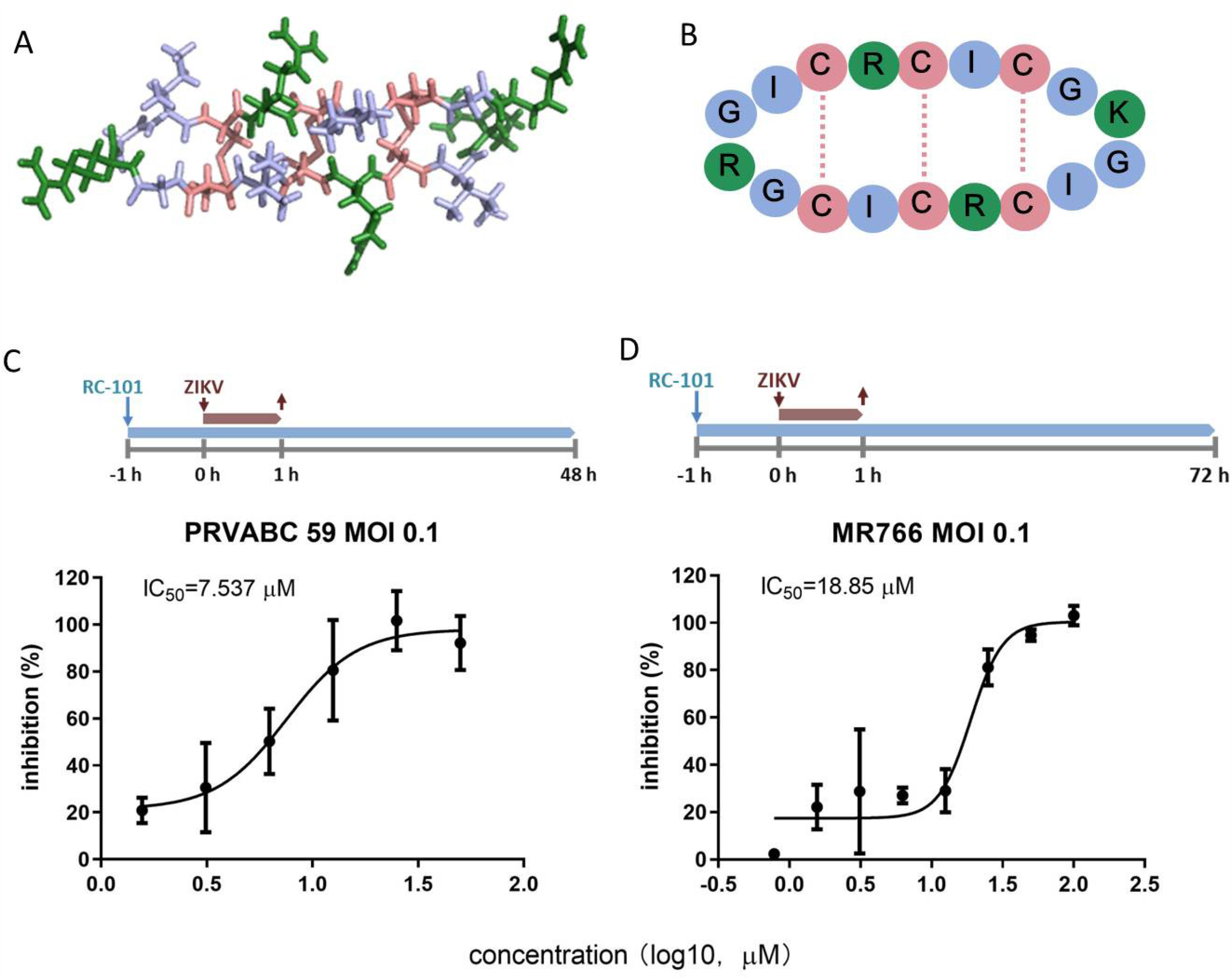
RC-101 inhibits Zika virus (ZIKV) infection. (A) Stick diagram of the crystal structure of RC-2 (PDB: 2LZI). (B) Schematic diagram of RC-101. Color in the schematic diagram correlates with those in the panel A. (C, D) RC-101 inhibits ZIKV strain PRVABC59 and strain MR766 infections. The experiments were carried out on Vero cells, and the experimental timeline is shown in C and D. After 48 or 72 h, the inhibitory effects were determined using an MTS assay.

## Results

### RC-101 inhibits ZIKV infection

To test the inhibitory effect of RC-101 against ZIKV infection, two strains were used to determine the 50% inhibitory concentration (IC_50_) of RC-101. Notably, the ZIKV PRVABC59 strain, belonging to the Asian-lineage ZIKV strains, contains one N-linked glycosylation site (N-X-S/T) at residue N154 of E, which is conserved among the flaviviruses, whereas the MR766 strain, belonging to the African lineage, lacks the glycosylation motif because of extensive passaging that leads to virus variants (26-31). A schematic of the assay is depicted in the upper panels of Fig. 1C and D. The incubation time of MR766 was 72 h, whereas that of PRVABC59 was 48 h, because the cytopathic effect of MR766 occurred 1 day after that of PRVABC59 with the same multiplicity of infection (MOI). As shown in Fig. 1C and D, RC-101 effectively blocked both ZIKV strain infections with IC_50_ values of 7.537 µM for PRVABC59 and 18.85 µM for MR766.

### RC-101 inhibits ZIKV infection at both the entry and replication steps

To test whether RC-101 blocked the entry step or the replication step, a time-of-addition experiment was performed (Fig. 2A). As shown in Fig. 2B and C, no suppression of viral titers was observed in the *pre-* or the *virucidal* treatment groups, indicating that RC-101 does not inhibit ZIKV infection either by blocking the cellular receptors that prevent virus binding or by inactivating the virus directly. However, RC-101 exerted significant inhibitory effects when its addition was synchronized with the virus in the *during* manner. Moreover, RC-101 inhibited MR766 strain infection when it was added 1 h post-infection. These results suggested that viral entry and replication are the stages at which RC-101 shows inhibitory activity.

**Fig. 2.**
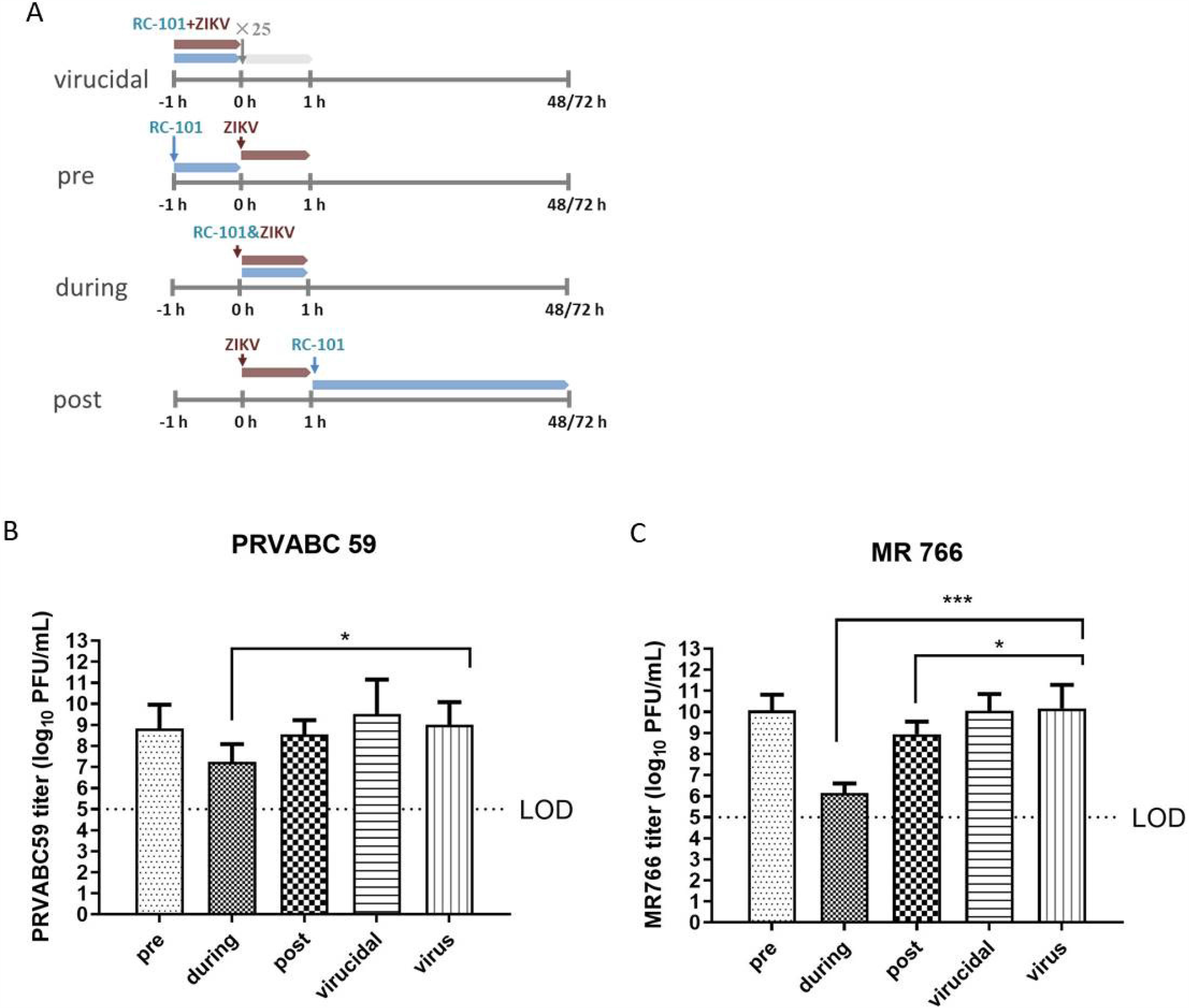
Time-of-addition analysis of the antiviral activity of the RC-101. (A) Schematic illustration of time-of-addition experiment. Vero cells were infected with Zika virus (ZIKV) PRVABC59 (B) and MR766 (C) (MOI: 0.1) for 1 h. RC-101 (40 µM) was introduced at different time points, designated as virucidal, pretreatment (pre), during treatment (during), or post-treatment (post). The inhibitory effect of the drugs in each group was determined by plaque assays.

To confirm the inhibitory effect on viral replication, we investigated the effects of RC-101 on ZIKV replicon. As shown in Fig. 3, RC-101 showed little effect on the initial translation of replicon RNA (32, 33) (Fig. 3A), whereas an appreciable reduction in the luciferase signal was observed at 48 h post-electroporation (Fig. 3B). This confirmed that RC-101 has an inhibitory effect on the ZIKV replication state.

**Fig. 3.**
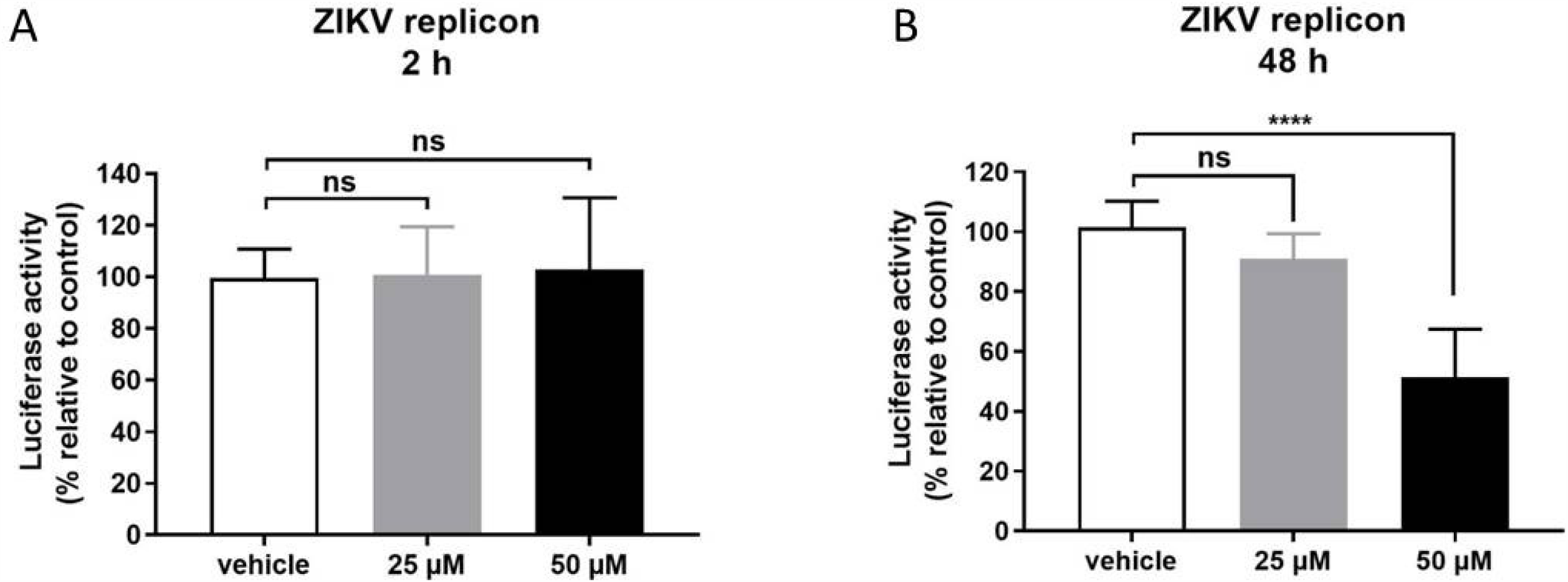
RC-101 inhibits Zika virus (ZIKV) replicon activity. (A, B) BHK-21 cells transfected with the ZIKV replicon were treated with RC-101 and luciferase activities were determined at 2 h (B) and 48 h (C).

### RC-101 inhibits NS2B-NS3 serine protease activity

To investigate the potential viral target of RC-101, we tested the inhibitory effect of RC-101 on ZIKV NS2B-NS3 protease activity. It has been reported that RC-1, which possesses the same residue sequence as RC-101, except for one lysine (K) instead of arginine (R) in RC-101, might dock at the NS2B and NS3 interface and thus inhibit DENV-2 replication by interfering with the activity of the NS2B-NS3 serine protease (20). Considering the sequence and structural conservation of flavivirus NS proteins, we reasoned that RC-101 might have a similar effect on the ZIKV NS2B-NS3 protease. To test this hypothesis, we first produced NS2B-NS3pro in *Escherichia coli* as a single-chain peptide (20, 34, 35). Protease activity was assessed using a fluorogenic peptide as a substrate at 37 °C for 30 min. As shown in Fig. 4A, the Michaelis-Menten constant (*Km*) value was 11.77 µM, indicating that the enzyme kinetic assay was robust and suitable to investigate the inhibitory effect. As shown in Fig. 4B, RC-101 effectively inhibited NS2B-NS3 protease activity with an IC_50_ of 7.20 µM, indicating that this protease serves as a viral target of RC-101.

**Fig. 4.**
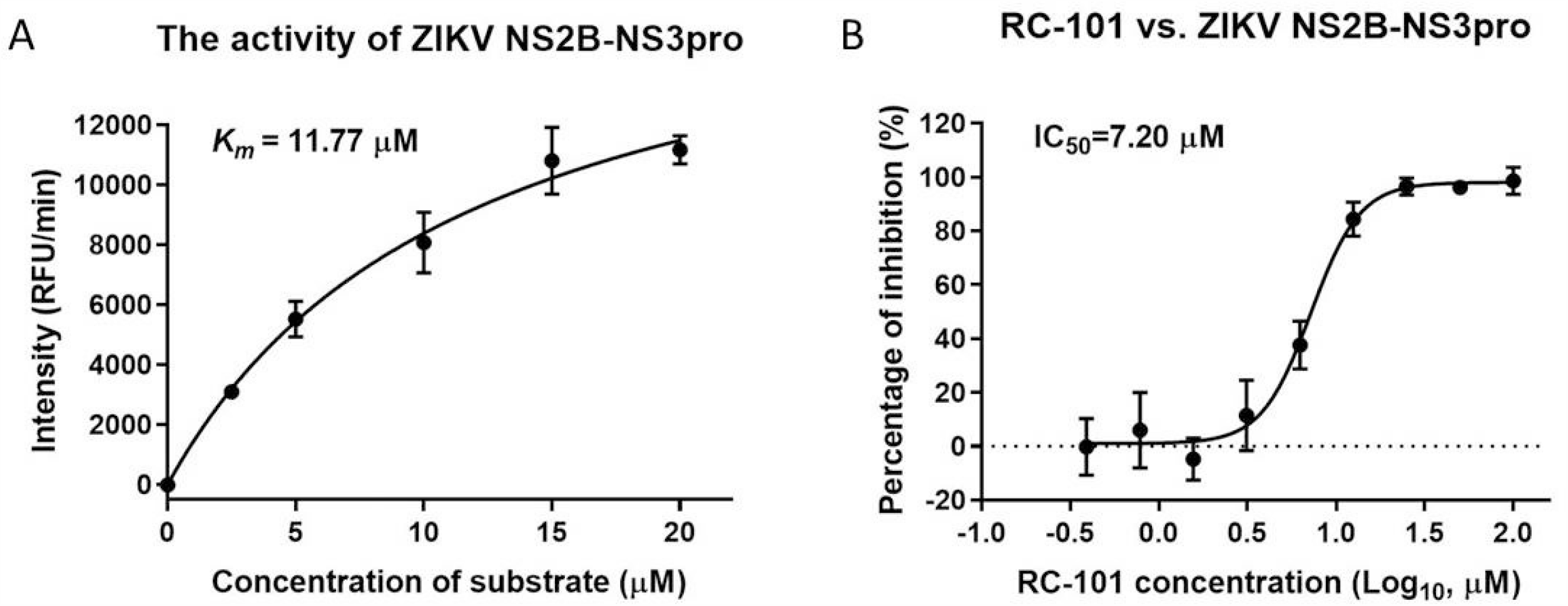
RC-101 inhibits the NS2B-NS3 serine protease activity. (A) Enzyme kinetic assay of NS2B-NS3pro activity. The fluorogenic substrate peptide (Boc-Gly-Arg-Arg-AMC) was serially diluted to assess the activity of Zika virus (ZIKV) protease. The relative fluorescence units (RFUs) were measured using an EnSpire multimode plate reader with the emission at 440 nm upon excitation at 350 nm. (B) The inhibitory effect of RC-101 against the activity of ZIKV NS2B-NS3pro. The reaction mixtures of NS2B-NS3pro (100 µl) consisted of 12 μM substrate peptide, 1.25 μM of NS2B-NS3pro, and RC-101 of varying concentrations with a buffer comprised of 200 mM tris-HCl (pH 8.5), and this was incubated at 37 °C for 30 min.

### RC-101 inhibits flavivirus entry by targeting the DE loop of E glycoprotein

As RC-101 was found to inhibit ZIKV infection both at the entry and replication stages (Fig. 2), we further investigated the mechanism underlying the inhibitory effect on the entry stage. As previously mentioned, RC has been reported to inhibit different types of enveloped viruses by binding to the negatively charged glycan chains on the surface of the glycoprotein, thus blocking virus entry (22-24). However, flaviviruses contain only one glycosylation motif on the E glycoprotein, but this the number is not absolutely conserved, as DENV has two glycosylation motifs, whereas some African-linage ZIKV strains have no glycan chain on the surface (26-31, 36-38). As shown in Fig. 1, RC-101 exerted similar inhibitory effects on both the ZIKV Asian strain PRVABC59 (one glycan) and the African strain MR766 (no glycan), suggesting that glycan might not be the target of RC-101. As RC-101 could block ZIKV infection at the entry stage (Fig. 2), we further investigated its effect on the E protein.

In our previously published work, we constructed a series of JEV variants with mutations in the receptor-binding motif or in amino acids critical for membrane fusion on the E protein (6). Considering the relative conservation of the sequence and structure of flavivirus E proteins, we used the constructed JEV variants to investigate the potential target of RC-101. Among the selected variants, the N154A and DE mutants were found to impair receptor binding by the virus, H144A and H319A abrogated the interaction between DI and DIII, and Q258A and T410A resulted in failure of the E protein to re-fold to form its post-fusion conformation (6). Notably, these six tested sites were conserved between JEV and ZIKV (Fig. 5). The investigation was conducted using the “during” manner (Fig. 6A). As shown in Fig. 6B and C, RC-101 at 50 µM, corresponding to the approximate IC_98_ against ZIKV (Fig. 1), robustly inhibited JEV infection, which made the prM band hardly detectable, and the viral titers decreased by approximately 3 log units. Similarly, RC-101 inhibited infections by viruses harboring N154A and H144A, suggesting that neither N154 nor H144 is the target of RC-101. Of note, the outcome indicating that abolishing the glycosylation motif (N154A) resulted in retained sensitivity to RC-101 was in line with the notion that differences in the number of glycan chains in different strains have little effect on RC-101 inhibition (Fig. 1). This further confirmed that RC-101 has a unique anti-flavivirus mechanism, which is unlike the effects on other enveloped viruses. Notably, as shown in Fig. 6B and C, the Q258A mutant likely had increased sensitivity to RC-101, whereas H319A resulted in resistance to RC-101 at the protein level and in the low MOI infection assay. Among the six tested mutants, the DE mutant and T410A showed robust resistance to RC-101 in all assays, indicating that these two mutants do confer resistance and might serve as the viral glycoprotein target(s) of RC-101. As T410 is located in the stem region of the E protein, buried by the compacted E dimer and hardly accessible in the prefusion conformation, the DE mutant was selected for further investigation of the binding affinity to RC-101.

**Fig. 5.**
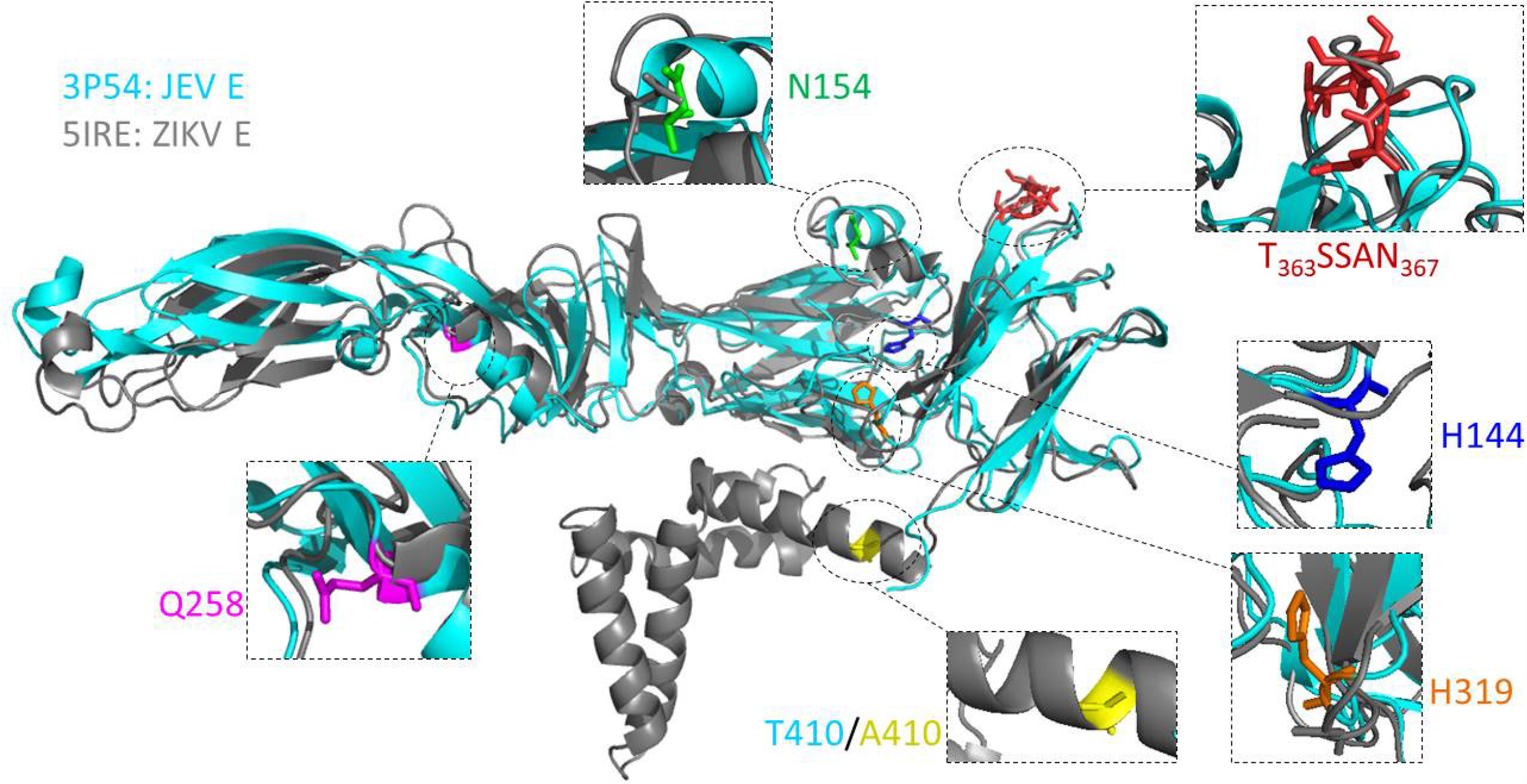
The potential viral target of RC-101 on flavivirus E protein. Side view of monomer prefusion Japanese encephalitis virus (JEV) E protein ectodomain conformation (cyan, PDB: 3P54) in alignment with the full-length Zika virus (ZIKV) E protein (gray, PDB: 5IRE). The potential targets tested in this study were enlarged and highlighted by colors.

**Fig. 6.**
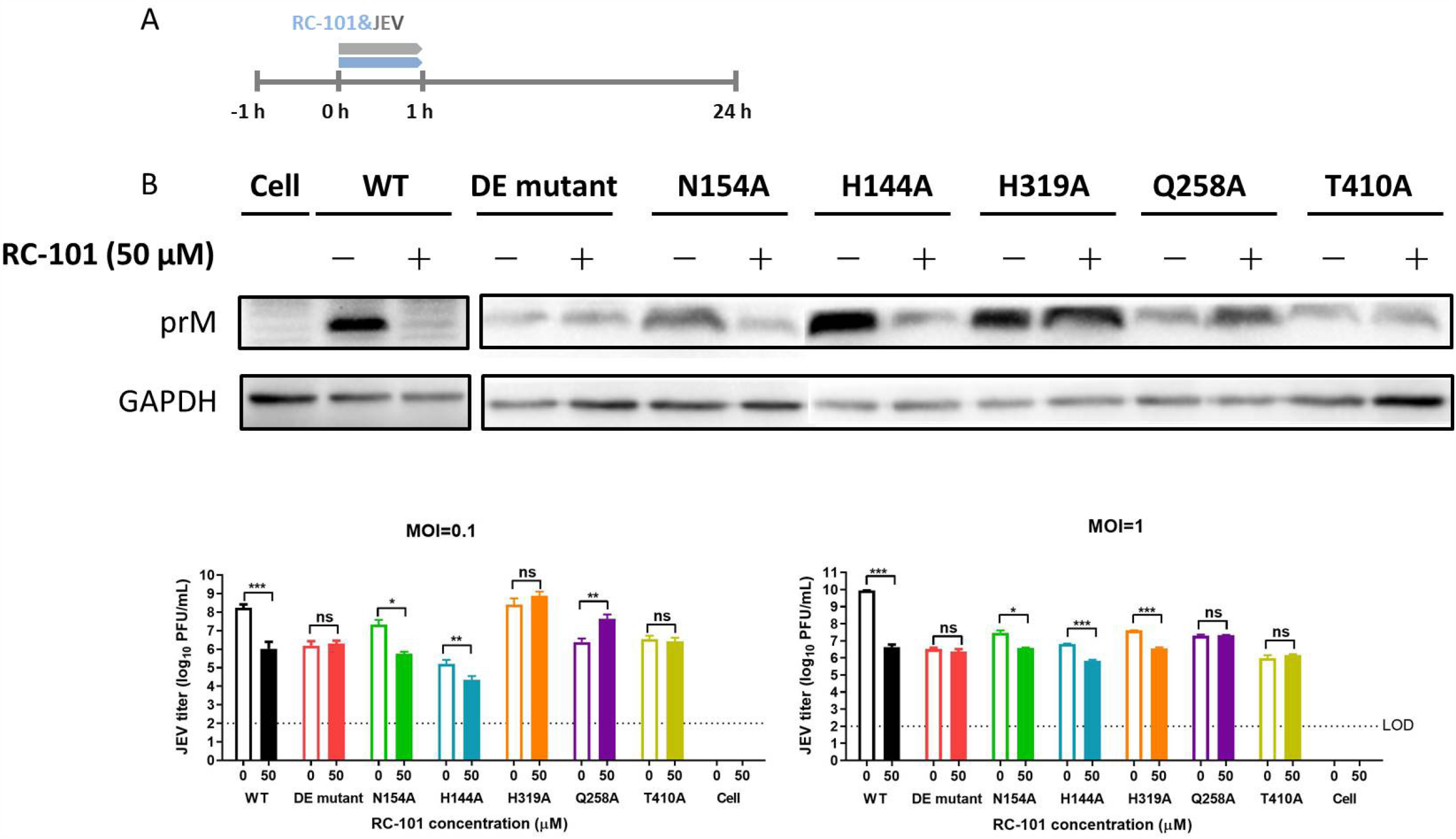
Sensitivity/resistance of the mutant viruses to RC-101. (A) Timeline of the assay. (B) BHK-21 cell lysates were analyzed by western blotting at 24 h post-infection, and rabbit prM antiserum, as well as the anti-GAPDH mouse monoclonal antibody, were used as primary antibodies. (C) The viral titers were tested by plaque forming assays using BHK-21 cells. Data are represented as the means ± SDs from 4–6 independent experiments. ***, *P* < 0.001; **, *P* < 0.01; *, *P* < 0.05.

### DE loop mutant decreases binding affinity to RC-101

To test the possibility that the DE loop is the target of RC-101, and to test whether the DE mutant would disrupt the binding of RC-101 to DIII, the binding affinities of WT and the DE mutant DIII to RC-101 were examined by biolayer interferometry. The interactions between DIII and RC-101 were calculated using a 1:1 binding model at three different concentrations (Fig. 7). The results showed that RC-101 bound to WT DIII with a kinetic association (*K*_a_) of 1.46 × 10^4^ M^−1^ s^−1^, kinetic dissociation (*K*_d_) of 1.18 × 10^−4^ s^−1^, and *K*_D_ of 8.10 × 10^−9^ M, indicating that RC-101 has high affinity for DIII. The binding affinity of RC-101 to the DE mutant was decreased by one order of magnitude, to a *K*_D_ with 2.37 × 10^−8^ M, which suggested that the DE loop might be the binding site of RC-101 and that the DE mutant would disrupt this interaction.

**Fig. 7.**
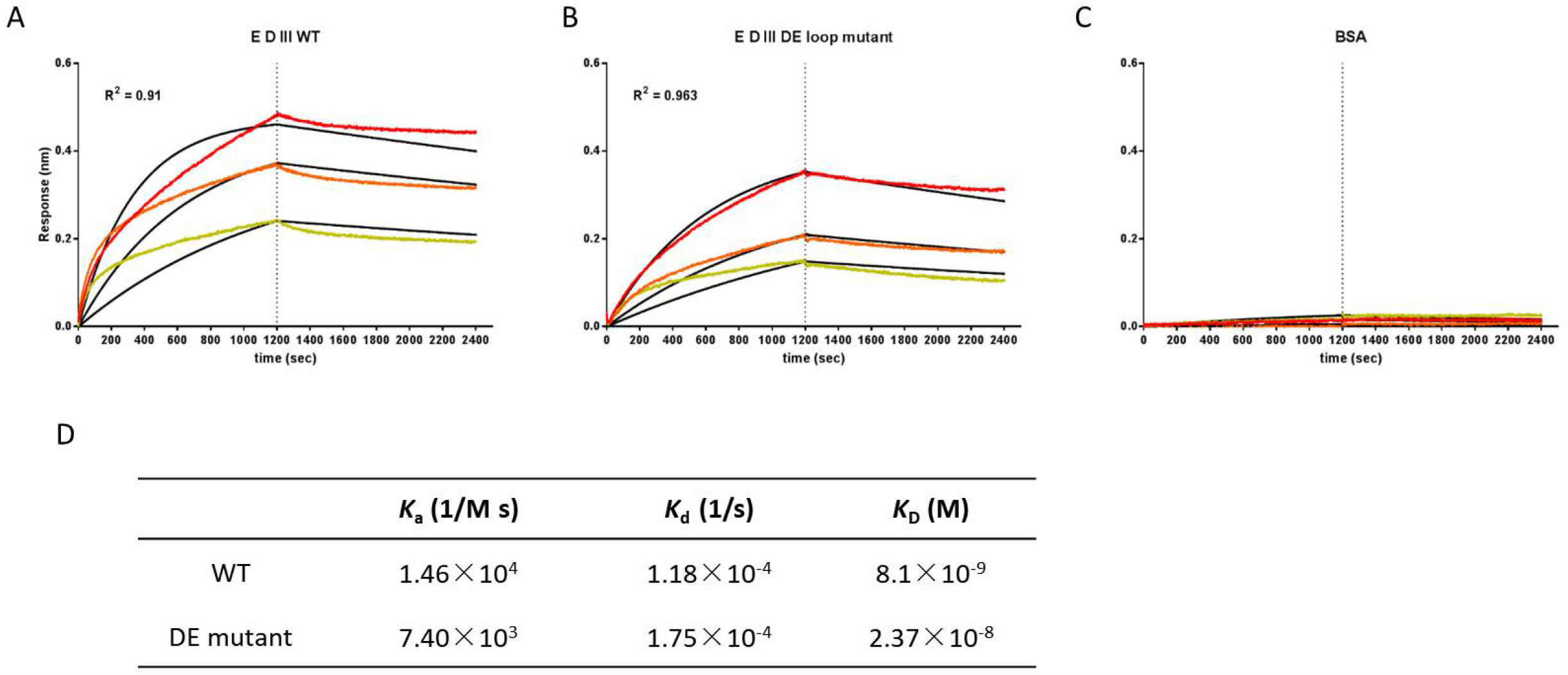
A DE loop mutation decreases the binding affinity of RC-101 to E protein domain III (DIII). WT DIII (A), DE loop mutant DIII (B), and BSA (C) were immobilized onto biosensors. The binding of RC-101 was assessed at 200 nM (red), 100 nM (orange), and 50 nM (yellow), and the global fit curves are shown as black lines. The vertical dashed lines indicate the transition between association and dissociation phases. (D) The binding affinities of WT and DE loop DIII to RC-101.

## Discussion

Although RC has been reported to have inhibitory effects against different kinds of viruses with various antiviral mechanisms, few studies have investigated its effect on flaviviruses. In this study, we evaluated the antiviral effects of RC-101 against flaviviruses and elucidate the mechanism of action. As the analogue RC-1 has been reported to inhibit DENV NS2B-NS3 protease and viral replication, we first tested whether RC-101 could extend its antiviral spectrum to other flaviviruses. As a result, RC-101 was found to inhibit infections by different strains of ZIKV, as well as JEV. Further, results suggest that the NS2B-NS3 protease might serve as one of the viral targets since RC-101 could block the serine protease activity of NS2B-NS3. The NS3 proteolytic domain forms a substrate-binding pocket with a catalytic triad, conserved in flaviviruses, of His-Asp-Ser (Fig. 8A). In an attempt to dock the analogue RC-2 (PDB: 2LZI) (39) with ZIKV NS3 (PDB: 5ZMS) (40), we found that glycine in RC-2 might interact with histidine (H1553) and serine (S1673) in the catalytic triad (Fig. 8B). RC-101 might thus inhibit NS2B-NS3 protease activity by competitively blocking the catalytic motif and thus preventing substrate binding. Meanwhile, as a cationic peptide, RC-101 might directly interact with the negatively charged NS2B and thus prevent the binding of NS2B and NS3 (20, 41).

**Fig. 8.**
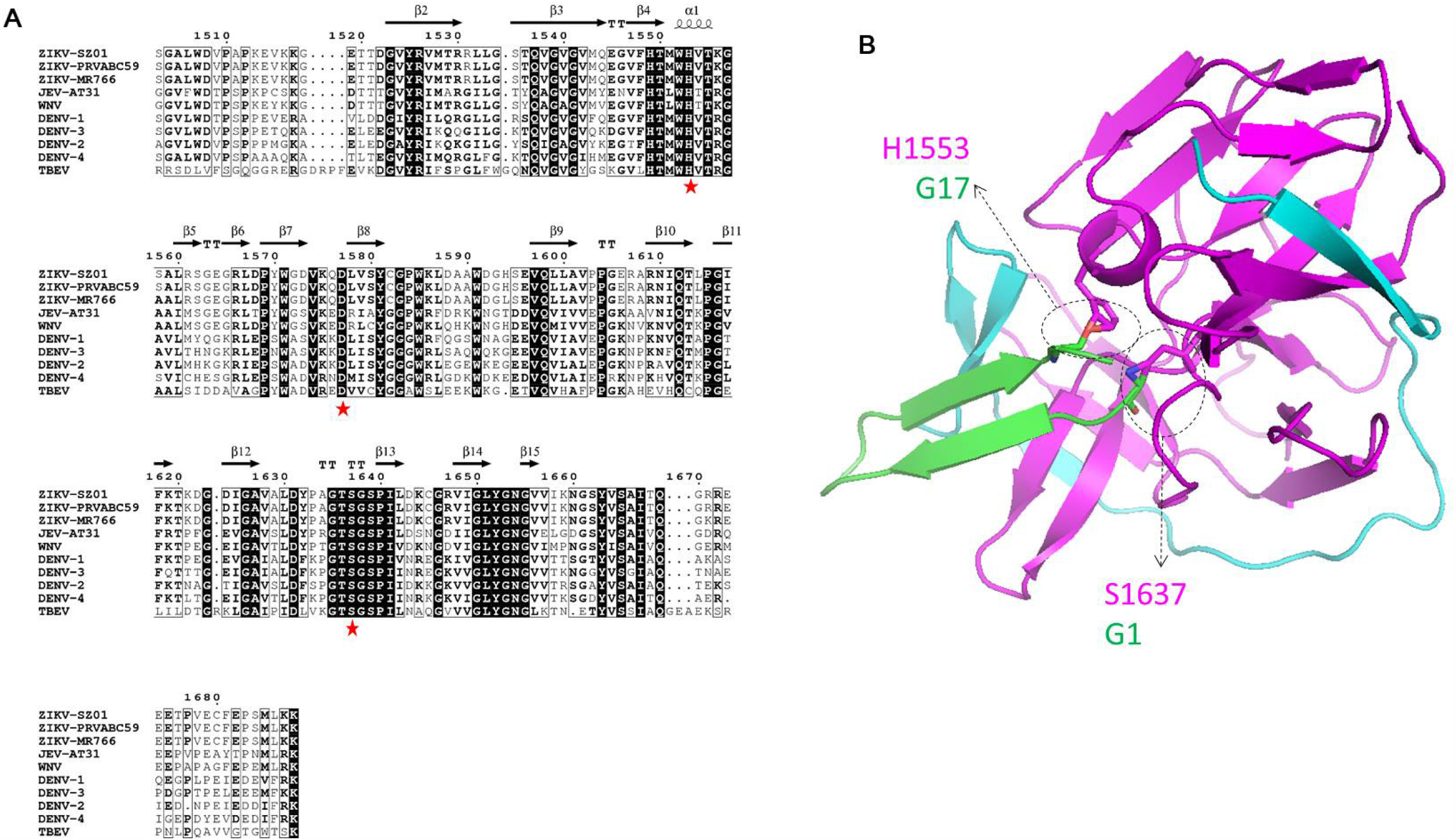
Docking of the NS2B-NS3/RC-2 complex. (A) Sequence alignment of the flavivirus NS3 N-terminal domain (1503–1688). Secondary structure elements were graphically represented by ESPript (51) (http://espript.ibcp.fr). The secondary structure observed with Zika virus (ZIKV) NS2B-NS3 protease (PDB: 5GXJ) is indicated above the sequence. The catalytic triad residues are indicated by a red asterisk. The relevant sequence accession numbers are as follows: ZIKV (strain SZ01,Genbank: KU866423.2), ZIKV (strain PRVABC59, MK713748.1), ZIKV (strain MR766, AY632535.2), Japanese encephalitis virus (JEV; strain AT31, AB196923.1), West Nile virus (WNV; NC_001563.2), dengue virus (DENV)-1 (AY145122.1), DENV-2 (NC_001474.2), DENV-3 (MN227700.1), DENV-4 (KY924607.1), Tick-borne encephalitis virus (MT311860.1) (B) The ribbon diagram of the NS2B-NS3/RC-2 complex. The crystal structure of RC-2 (PDB 2ZLI) and ZIKV NS3 (PDB: 5ZMS) was used to build the complex using the ZDOCK 3.0.2 program. NS2B, NS3, and RC-2 are colored as cyan, magenta, and green, respectively. The supposed interacting residues between NS3 and RC-2 are shown as sticks.

As mentioned previously herein, RC has been extensively reported to inhibit enveloped viruses by targeting the negative glycan shield on the surface of the virus, thus blocking the initial entry of the virus into host cells (22-24). As the only glycan chain in the E protein of ZIKV PRVABC59 strain and JEV, the glycan linked to the N_154_YS glycosylation motif has been reported to interact with DC-SIGN, which is a candidate flavivirus receptor (42). Intriguingly, the N154A mutation had no impact on the sensitivity or resistance of JEV to RC-101. A possible explanation for this phenomenon is that RC could easily bind the dense glycan shield of gp120 and HA of HIV and IAV, respectively, but for the flavivirus, RC might pass through the unique glycan and interact with the E protein directly. The DE loop, which is the relatively higher tip of the E protein (Fig. 5), might serve as the viral target of RC. Although peptides derived from the DE loop were previously found to prevent JEV infection by interfering with virus attachment to BHK-21 cells (43), the DE loop is not the only or major receptor binding motif for JEV entry into different types of cells (6). Further studies should focus on whether RC-101 could inhibit flavivirus infection of different kinds of cells and whether the DE mutant confers resistance to RC-101 in other hosts and tissues.

Currently, there are no effective drugs approved for the treatment of flavivirus infection. Fortunately, several peptide inhibitors, derived from the E protein or targeting the E protein, have been used to successfully block flavivirus infection *in vitro* and *in vivo* (7, 9, 12, 44). As the flavivirus E protein has a highly conserved sequence and conformation, peptide inhibitors could be used for the treatment of emerging flavivirus infections or severe cases. In addition, peptide inhibitors have many advantages, such as high biocompatibility, a low frequency of selecting resistant mutants, the ability to synergize with conventional drugs, and activity towards multi-drug resistant virus strains (45). The cyclic peptide RC, with a unique structure that provides long-lasting protection against viral infection, is a potential candidate for the development of a successful drug to treat flaviviruses and other infectious diseases.

## Materials and Methods

### Cells, viruses, and RC-101

Vero and BHK-21 cells were maintained in Dulbecco’s modified Eagle’s medium and minimum essential medium containing 10% fetal bovine serum, respectively. The ZIKV strains PRVABC59 (GenBank accession no. KX377337.1) and MR-776 (GenBank accession no. KX377335.1) were kindly provided by Jean K Lim (Icahn School of Medicine at Mount Sinai, NY). The genome sequence of ZIKV strain SZ-WIV001 (GenBank accession no.KU963796) was used as the template for the construction of the ZIKV replicon (46). JEV AT31 was generated using the infectious clones of pMWJEAT AT31 (kindly provided by T. Wakita, Tokyo Metropolitan Institute for Neuroscience) as previously described (47). The JEV variants, including the DE mutant, N154A, H144A, H319A, Q258A, and T410A, were constructed and preserved at −80 °C in our laboratory (6). RC-101was was synthesized by solid-phase synthesis and purified by reversed-phase HPLC to homogeneity (98% purity) (21).

### Antiviral effects of RC-101

Vero cells in 96-well plates were infected with ZIKV PRVABC59 and MR-766 at the indicated MOI in the presence of RC-101 at different concentrations for 48 and 72 h, respectively. The antiviral effects were evaluated by an MTS assay.

Time-of-addition assay. Vero cells were infected with ZIKV (MOI, 0.1) for 1 h (0–1 h). RC-101 was incubated with the cells for 1 h before infection (−1 to 0 h), during infection (0 to 1 h), and for 47 or 71 h post-infection (Fig. 2A). To exclude a possible direct inactivating effect of RC-101, ZIKV was incubated with RC-101 at 37 °C for 1 h, and the mixtures were diluted 25-fold to infect Vero cells. To confirm the inhibitory effect of RC-101 against ZIKV replication, BHK-21 cells were electroporated with the ZIKV replicon (SZ-WIV001; Genbank No: KU963796) and then incubated with RC-101. *Renilla* luciferase activity in the cell lysates was measured using the Rluc system (Promega, Madison, WI, USA) (48).

### Proteolytic activity of NS2B-NS3 protease

To produce NS2B-GGGGSGGGG-NS3 protein, the ZIKV replicon was used as the template, and the NS2B fragments were amplified by PCR using primer pairs (forward: 5′-TTAAGAAGGAGATATA<u>CCATGG</u>GCGTGGACATGTACATTGAAAGAG-3′; reverse: 5′-CACCACT*TCCACCTCCACCCGATCCACCTCCACC*GATCTCTCTCATGGGGGG ACC-3′), and NS3 was also amplified using primer pairs (forward: 5′-GAGATC*GGTGGAGGTGGATCGGGTGGAGGTGGA*AGTGGTGCTCTATGGGAT GTGC-3′, reverse: 5′-CTCAGTGGTGGTGGTGGTGGTG<u>CTCGAG</u>CTTCTTCAGCATCGAAGGCTC GAAG-3′) (20). The PCR products were cloned into pET28a using infusion PCR (Novagen, Darmstadt, Germany). The recombinant vector was transformed into *E. coli* BL21(DE3), and the cell lysates were loaded onto a nickel column. The protein was eluted with a gradient concentration of imidazole buffer (50 mM tris-HCl, 30 mM NaCl, 50–500 mM imidazole, pH 7.0) (35).

The proteolytic activity of NS2B-NS3pro was measured using a fluorescence resonance energy transfer-based assay with a fluorogenic peptide substrate (Boc-Gly-Arg-Arg-AMC, No: I-1565, Bachem) as the substrate. The relative fluorescence units were measured using an EnSpire multimode plate reader with the emission at 440 nm upon excitation at 350 nm. The kinetic parameter of NS2B-NS3pro was obtained using substrate from 2.5 to 20 µM in the fluorescent assay after a 30-min incubation at 37 °C (20, 49). The *Km* was calculated from the enzyme kinetics-velocity as a function of substrate model using GraphPad Prism 8.0. The inhibitory effects of RC-101 against protease activities was assessed at 37 °C for 30 min, with mixtures of 100 μl consisting of 12 μM fluorogenic peptide substrate, 1.25 μM of NS2B-NS3pro, and RC-101 ranging from 0 to 100 μM, buffered at pH 8.5 with 200 mM tris–HCl. The IC_50_ value of RC-101 was evaluated using the non-linear regression model in GraphPad Prism 8.0.

### Expression of WT and DE mutant DIII

The WT DIII expression vector was constructed using pET-22b(+) and preserved in our laboratory (7). The DE mutant was constructed using the East Mutagenesis System Kit (TransGen Biotech, China) with the following primer pairs (forward: 5′-CAGTGAACCCCTTCGTCGCGGCGGCGGCGGCGGCGTCAAAGGTGC-3′; reverse: 5′-CGCCGCCGCCGCCGCCGCGACGAAGGGGTTCACTGTCACCAGCCG-3′) (6). WT DIII was expressed using *E. coli* BL21 (DE3); the supernatant of the bacterial pellets was loaded onto a nickel column, and the bound protein was eluted with a gradient concentration of imidazole buffer. DE mutant DIII, expressed as inclusion bodies, was solubilized in 8 M urea (50 mM tris-HCl, 100 mM NaCl, 1mM DTT, 0.1% SDS, 8 M urea, pH 7.4). Refolding was carried out by titration dialysis at 4 °C against refolding buffer (50 mM tris-HCl, 100 mM NaCl, 0.1% SDS, 1 mM L(+)-arginine, 1 mM glutathione, 5% glycerine, pH 7.4) until the concentration of urea was < 2 M. Then, the supernatant was passed through a nickel column as described previously herein.

### Binding affinity assay

Real-time binding assays between RC-101 and WT or the DE mutant DIII were performed using biolayer interferometry on an Octet QK system (Fortebio, USA) according to previously reported methods (7). Binding kinetics were calculated using the Octet QK software package, which fit the observation to a 1:1 model to calculate the association and dissociation rate constants. Binding affinities were calculated as the *K*_d_ rate constant divided by the *K*_a_ rate constant.

### Docking of the NS2B-NS3/RC-2 complex

The crystal structures of RC-2 (PDB 2ZLI) and ZIKV NS3 (PDB: 5ZMS) were used to build the complex using the ZDOCK 3.0.2 program (http://zdock.umassmed.edu) (50). The resulting model was represented by PyMOL.

## ACKNOWLEDGEMENTS

We thank the Center for Instrumental Analysis and Metrology, and Core Facility and Technical Support, Wuhan Institute of Virology, for providing technical assistance. This work was supported by the National Key Research and Development Program of China (2018YFA0507204), the National Natural Sciences Foundation of China (31670165), Wuhan National Biosafety Laboratory, Chinese Academy of Sciences Advanced Customer Cultivation Project (2019ACCP-MS03), the Open Research Fund Program of the State Key Laboratory of Virology of China (2018IOV001).

## References

1. Mackenzie JS, Gubler DJ, Petersen LR. 2004. Emerging flaviviruses: the spread and resurgence of Japanese encephalitis, West Nile and dengue viruses. Nat Med 10:S98–109.

2. Unni SK, Ruzek D, Chhatbar C, Mishra R, Johri MK, Singh SK. 2011. Japanese encephalitis virus: from genome to infectome. Microbes Infect 13:312–21.

3. Rey FA, Heinz FX, Mandl C, Kunz C, Harrison SC. 1995. The envelope glycoprotein from tick-borne encephalitis virus at 2 A resolution. Nature 375:291–8.

4. Zhao H, Fernandez E, Dowd KA, Speer SD, Platt DJ, Gorman MJ, Govero J, Nelson CA, Pierson TC, Diamond MS, Fremont DH. 2016. Structural Basis of Zika Virus-Specific Antibody Protection. Cell 166:1016–27.

5. Luca VC, AbiMansour J, Nelson CA, Fremont DH. 2012. Crystal structure of the Japanese encephalitis virus envelope protein. J Virol 86:2337–46.

6. Liu H, Liu Y, Wang S, Zhang Y, Zu X, Zhou Z, Zhang B, Xiao G. 2015. Structure-based mutational analysis of several sites in the E protein: implications for understanding the entry mechanism of Japanese encephalitis virus. J Virol 89:5668–86.

7. Zu X, Liu Y, Wang S, Jin R, Zhou Z, Liu H, Gong R, Xiao G, Wang W. 2014. Peptide inhibitor of Japanese encephalitis virus infection targeting envelope protein domain III. Antiviral Res 104:7–14.

8. Lee E, Weir RC, Dalgarno L. 1997. Changes in the dengue virus major envelope protein on passaging and their localization on the three-dimensional structure of the protein. Virology 232:281–90.

9. Chen L, Liu Y, Wang S, Sun J, Wang P, Xin Q, Zhang L, Xiao G, Wang W. 2017. Antiviral activity of peptide inhibitors derived from the protein E stem against Japanese encephalitis and Zika viruses. Antiviral Res 141:140–149.

10. Bressanelli S, Stiasny K, Allison SL, Stura EA, Duquerroy S, Lescar J, Heinz FX, Rey FA. 2004. Structure of a flavivirus envelope glycoprotein in its low-pH-induced membrane fusion conformation. EMBO J 23:728–38.

11. Modis Y, Ogata S, Clements D, Harrison SC. 2004. Structure of the dengue virus envelope protein after membrane fusion. Nature 427:313–9.

12. Yu Y, Deng YQ, Zou P, Wang Q, Dai Y, Yu F, Du L, Zhang NN, Tian M, Hao JN, Meng Y, Li Y, Zhou X, Fuk-Woo Chan J, Yuen KY, Qin CF, Jiang S, Lu L. 2017. A peptide-based viral inactivator inhibits Zika virus infection in pregnant mice and fetuses. Nat Commun 8:15672.

13. Wang Q, Yang H, Liu X, Dai L, Ma T, Qi J, Wong G, Peng R, Liu S, Li J, Li S, Song J, Liu J, He J, Yuan H, Xiong Y, Liao Y, Li J, Yang J, Tong Z, Griffin BD, Bi Y, Liang M, Xu X, Qin C, Cheng G, Zhang X, Wang P, Qiu X, Kobinger G, Shi Y, Yan J, Gao GF. 2016. Molecular determinants of human neutralizing antibodies isolated from a patient infected with Zika virus. Sci Transl Med 8:369ra179.

14. Luo D, Vasudevan SG, Lescar J. 2015. The flavivirus NS2B-NS3 protease-helicase as a target for antiviral drug development. Antiviral Res 118:148–58.

15. Sampath A, Padmanabhan R. 2009. Molecular targets for flavivirus drug discovery. Antiviral Res 81:6–15.

16. Arnett E, Lehrer RI, Pratikhya P, Lu W, Seveau S. 2011. Defensins enable macrophages to inhibit the intracellular proliferation of Listeria monocytogenes. Cell Microbiol 13:635–51.

17. Leonova L, Kokryakov VN, Aleshina G, Hong T, Nguyen T, Zhao C, Waring AJ, Lehrer RI. 2001. Circular minidefensins and posttranslational generation of molecular diversity. J Leukoc Biol 70:461–4.

18. Tang YQ, Yuan J, Osapay G, Osapay K, Tran D, Miller CJ, Ouellette AJ, Selsted ME. 1999. A cyclic antimicrobial peptide produced in primate leukocytes by the ligation of two truncated alpha-defensins. Science 286:498–502.

19. Tran D, Tran PA, Tang YQ, Yuan J, Cole T, Selsted ME. 2002. Homodimeric theta-defensins from rhesus macaque leukocytes: isolation, synthesis, antimicrobial activities, and bacterial binding properties of the cyclic peptides. J Biol Chem 277:3079–84.

20. Rothan HA, Han HC, Ramasamy TS, Othman S, Rahman NA, Yusof R. 2012. Inhibition of dengue NS2B-NS3 protease and viral replication in Vero cells by recombinant retrocyclin-1. BMC Infect Dis 12:314.

21. Prantner D, Shirey KA, Lai W, Lu W, Cole AM, Vogel SN, Garzino-Demo A. 2017. The theta-defensin retrocyclin 101 inhibits TLR4-and TLR2-dependent signaling and protects mice against influenza infection. J Leukoc Biol 102:1103–1113.

22. Wang W, Cole AM, Hong T, Waring AJ, Lehrer RI. 2003. Retrocyclin, an antiretroviral theta-defensin, is a lectin. J Immunol 170:4708–16.

23. Leikina E, Delanoe-Ayari H, Melikov K, Cho MS, Chen A, Waring AJ, Wang W, Xie Y, Loo JA, Lehrer RI, Chernomordik LV. 2005. Carbohydrate-binding molecules inhibit viral fusion and entry by crosslinking membrane glycoproteins. Nat Immunol 6:995–1001.

24. Yasin B, Wang W, Pang M, Cheshenko N, Hong T, Waring AJ, Herold BC, Wagar EA, Lehrer RI. 2004. Theta defensins protect cells from infection by herpes simplex virus by inhibiting viral adhesion and entry. J Virol 78:5147–56.

25. Carbaugh DL, Lazear HM. 2020. Flavivirus Envelope Protein Glycosylation: Impacts on Viral Infection and Pathogenesis. J Virol 94.

26. Goo L, DeMaso CR, Pelc RS, Ledgerwood JE, Graham BS, Kuhn RJ, Pierson TC. 2018. The Zika virus envelope protein glycan loop regulates virion antigenicity. Virology 515:191–202.

27. Frumence E, Viranaicken W, Bos S, Alvarez-Martinez MT, Roche M, Arnaud JD, Gadea G, Despres P. 2019. A Chimeric Zika Virus between Viral Strains MR766 and BeH819015 Highlights a Role for E-glycan Loop in Antibody-mediated Virus Neutralization. Vaccines (Basel) 7.

28. Fontes-Garfias CR, Shan C, Luo H, Muruato AE, Medeiros DBA, Mays E, Xie X, Zou J, Roundy CM, Wakamiya M, Rossi SL, Wang T, Weaver SC, Shi PY. 2017. Functional Analysis of Glycosylation of Zika Virus Envelope Protein. Cell Rep 21:1180–1190.

29. Carbaugh DL, Baric RS, Lazear HM. 2019. Envelope Protein Glycosylation Mediates Zika Virus Pathogenesis. J Virol 93.

30. Beaver JT, Lelutiu N, Habib R, Skountzou I. 2018. Evolution of Two Major Zika Virus Lineages: Implications for Pathology, Immune Response, and Vaccine Development. Front Immunol 9:1640.

31. Annamalai AS, Pattnaik A, Sahoo BR, Muthukrishnan E, Natarajan SK, Steffen D, Vu HLX, Delhon G, Osorio FA, Petro TM, Xiang SH, Pattnaik AK. 2017. Zika Virus Encoding Nonglycosylated Envelope Protein Is Attenuated and Defective in Neuroinvasion. J Virol 91.

32. Puig-Basagoiti F, Deas TS, Ren P, Tilgner M, Ferguson DM, Shi PY. 2005. High-throughput assays using a luciferase-expressing replicon, virus-like particles, and full-length virus for West Nile virus drug discovery. Antimicrob Agents Chemother 49:4980–8.

33. Wang S, Liu H, Zu X, Liu Y, Chen L, Zhu X, Zhang L, Zhou Z, Xiao G, Wang W. 2016. The ubiquitin-proteasome system is essential for the productive entry of Japanese encephalitis virus. Virology 498:116–127.

34. Lei J, Hansen G, Nitsche C, Klein CD, Zhang L, Hilgenfeld R. 2016. Crystal structure of Zika virus NS2B-NS3 protease in complex with a boronate inhibitor. Science 353:503–5.

35. Lim HJ, Nguyen TT, Kim NM, Park JS, Jang TS, Kim D. 2017. Inhibitory effect of flavonoids against NS2B-NS3 protease of ZIKA virus and their structure activity relationship. Biotechnol Lett 39:415–421.

36. Chambers TJ, Hahn CS, Galler R, Rice CM. 1990. Flavivirus genome organization, expression, and replication. Annu Rev Microbiol 44:649–88.

37. Lee E, Leang SK, Davidson A, Lobigs M. 2010. Both E protein glycans adversely affect dengue virus infectivity but are beneficial for virion release. J Virol 84:5171–80.

38. Johnson AJ, Guirakhoo F, Roehrig JT. 1994. The envelope glycoproteins of dengue 1 and dengue 2 viruses grown in mosquito cells differ in their utilization of potential glycosylation sites. Virology 203:241–9.

39. Conibear AC, Rosengren KJ, Harvey PJ, Craik DJ. 2012. Structural characterization of the cyclic cystine ladder motif of theta-defensins. Biochemistry 51:9718–26.

40. Phoo WW, Zhang Z, Wirawan M, Chew EJC, Chew ABL, Kouretova J, Steinmetzer T, Luo D. 2018. Structures of Zika virus NS2B-NS3 protease in complex with peptidomimetic inhibitors. Antiviral Res 160:17–24.

41. Erbel P, Schiering N, D’Arcy A, Renatus M, Kroemer M, Lim SP, Yin Z, Keller TH, Vasudevan SG, Hommel U. 2006. Structural basis for the activation of flaviviral NS3 proteases from dengue and West Nile virus. Nat Struct Mol Biol 13:372–3.

42. Pokidysheva E, Zhang Y, Battisti AJ, Bator-Kelly CM, Chipman PR, Xiao C, Gregorio GG, Hendrickson WA, Kuhn RJ, Rossmann MG. 2006. Cryo-EM reconstruction of dengue virus in complex with the carbohydrate recognition domain of DC-SIGN. Cell 124:485–93.

43. Li C, Zhang LY, Sun MX, Li PP, Huang L, Wei JC, Yao YL, Isahg H, Chen PY, Mao X. 2012. Inhibition of Japanese encephalitis virus entry into the cells by the envelope glycoprotein domain III (EDIII) and the loop3 peptide derived from EDIII. Antiviral Res 94:179–83.

44. Schmidt AG, Yang PL, Harrison SC. 2010. Peptide inhibitors of flavivirus entry derived from the E protein stem. J Virol 84:12549–54.

45. Batoni G, Maisetta G, Brancatisano FL, Esin S, Campa M. 2011. Use of antimicrobial peptides against microbial biofilms: advantages and limits. Curr Med Chem 18:256–79.

46. Li JQ, Deng CL, Gu D, Li X, Shi L, He J, Zhang QY, Zhang B, Ye HQ. 2018. Development of a replicon cell line-based high throughput antiviral assay for screening inhibitors of Zika virus. Antiviral Res 150:148–154.

47. Li XD, Li XF, Ye HQ, Deng CL, Ye Q, Shan C, Shang BD, Xu LL, Li SH, Cao SB, Yuan ZM, Shi PY, Qin CF, Zhang B. 2014. Recovery of a chemically synthesized Japanese encephalitis virus reveals two critical adaptive mutations in NS2B and NS4A. J Gen Virol 95:806–15.

48. Guo J, Jia X, Liu Y, Wang S, Cao J, Zhang B, Xiao G, Wang W. 2020. Inhibition of Na(+)/K(+) ATPase blocks Zika virus infection in mice. Commun Biol 3:380.

49. Rothan HA, Abdulrahman AY, Sasikumer PG, Othman S, Rahman NA, Yusof R. 2012. Protegrin-1 inhibits dengue NS2B-NS3 serine protease and viral replication in MK2 cells. J Biomed Biotechnol 2012:251482.

50. Pierce BG, Wiehe K, Hwang H, Kim BH, Vreven T, Weng Z. 2014. ZDOCK server: interactive docking prediction of protein-protein complexes and symmetric multimers. Bioinformatics 30:1771–3.

51. Robert X, Gouet P. 2014. Deciphering key features in protein structures with the new ENDscript server. Nucleic Acids Res 42:W320–4.

